# Computational correction of off-targeting for CRISPR-Cas9 essentiality screens

**DOI:** 10.1101/809970

**Authors:** Alexendar R. Perez, Laura Sala, Richard K. Perez, Joana A. Vidigal

## Abstract

Off-target cleavage by Cas9 can confound measurements of cell proliferation/viability in CRISPR assays by eliciting a DNA-damage response that includes cell cycle arrest^1-3^. This gene-independent toxicity has been documented in large scale assays^2-4^ and shown to be a source of false-positives when libraries are populated by promiscuous guide RNAs (gRNAs)^7^. To address this, we developed CSC, a computational method to correct for the effect of specificity on gRNA depletion. We applied CSC to screening data from the Cancer Dependency Map and show that it significantly improves the specificity of CRISPR-Cas9 essentiality screens while preserving known gene essentialities even for genes targeted by highly promiscuous guides. We packaged CSC in a Python software to allow its seamless integration into current CRISPR analysis pipelines and improve the sensitivity of essentiality screens for repetitive genomic loci.

---

High-throughput loss-of-function screens can help catalog loci essential to cellular fitness^5-8^ and have been leveraged to systematically identify cancer vulnerabilities that can be exploited therapeutically^5^. The CRISPR-Cas9 genome editing system has become instrumental in these efforts, owing to the ease at which null alleles can be generated in a multiplex manner for both coding and non-coding regions.

Nevertheless, measurements of cell fitness in CRISPR genome-wide loss-of-function screens can be confounded by off-target cleavage because gRNAs that lead Cas9 to cleave multiple loci can trigger a DNA-damage response that includes cell cycle arrest^9^. The consequences of off-target cleavage to screen performance have been best characterized for gRNAs targeting amplified genomic regions^1, 10^ but have also been well described for promiscuous gRNAs within published genome-wide libraries both when they have perfect alignment or single mismatches to off-target sites^3^. While sophisticated computational strategies have been developed to correct for copy number effect on gRNA depletion^11, 12^, current pipelines handle unspecific gRNAs through the implementation of filters that remove guides suspected of off-target activity from further analysis^1, 6^. This strategy limits the information retrieved from unspecific libraries, and is incompatible with fitness screens to a large fraction of non-coding regulatory sequences^2^ since they are often repetitive in nature and therefore prone to be targeted by unspecific gRNAs^2, 13^. To address this, we developed CRISPR Specificity Correction (CSC), a computational approach to remove the confounding effect of off-targeting from high-throughput CRISPR fitness screens.

To systematically evaluate the effect of specificity on gRNA depletion we analyzed loss-of-function screens from the DepMap 19Q4 release of the public Avana dataset of the Cancer Dependency Map initiative^6, 11^. This dataset comprises screens performed in 26 distinct cellular lineages (**Supplementary Figure 1a**). We enumerated off-targets for each gRNA in the Avana library using GuideScan^13^, a retrieval-tree-based algorithm that outperforms short-read aligners in this task (**Supplementary Note 1**). Our estimates of potential off-target loci for Avana surpass those originally reported for this library^14^ as well as the estimates used by Project Achilles in the DepMap data processing pipeline^6^ (**Supplementary Table 1, Supplementary Figure 1b** and **1c**, **Supplementary Note 1**). To summarize the specificity of each gRNA in this library, we also computed GuideScan’s specificity score. This score aggregates Cutting Frequency Determination values (or CFD, describing the likelihood of an off-target being cut by Cas9 based on the number, position, and identity of mismatches to the gRNA)^14^ for all potential target sites enumerated by Guidescan^13^, so that the most specific targeting gRNAs receive a score of 1 and the most unspecific a score of 0. In agreement with previous studies^2, 3^, gRNAs with low specificities were on average more depleted from the population, often beyond the levels observed for gRNAs targeting known essential genes (**Figure 1a, left**). This observation held true even for guides with only mismatched off-targets (**Figure 1b**). Of note, distributions of gRNAs with scores above 0.16 became statistically indistinguishable from those of highly specific guides (Kolmogorov–Smirnov test, adjusted for multiple testing) suggesting that above this threshold off-targeting minimally influences the representation of gRNAs in the library.

**Figure 1.**
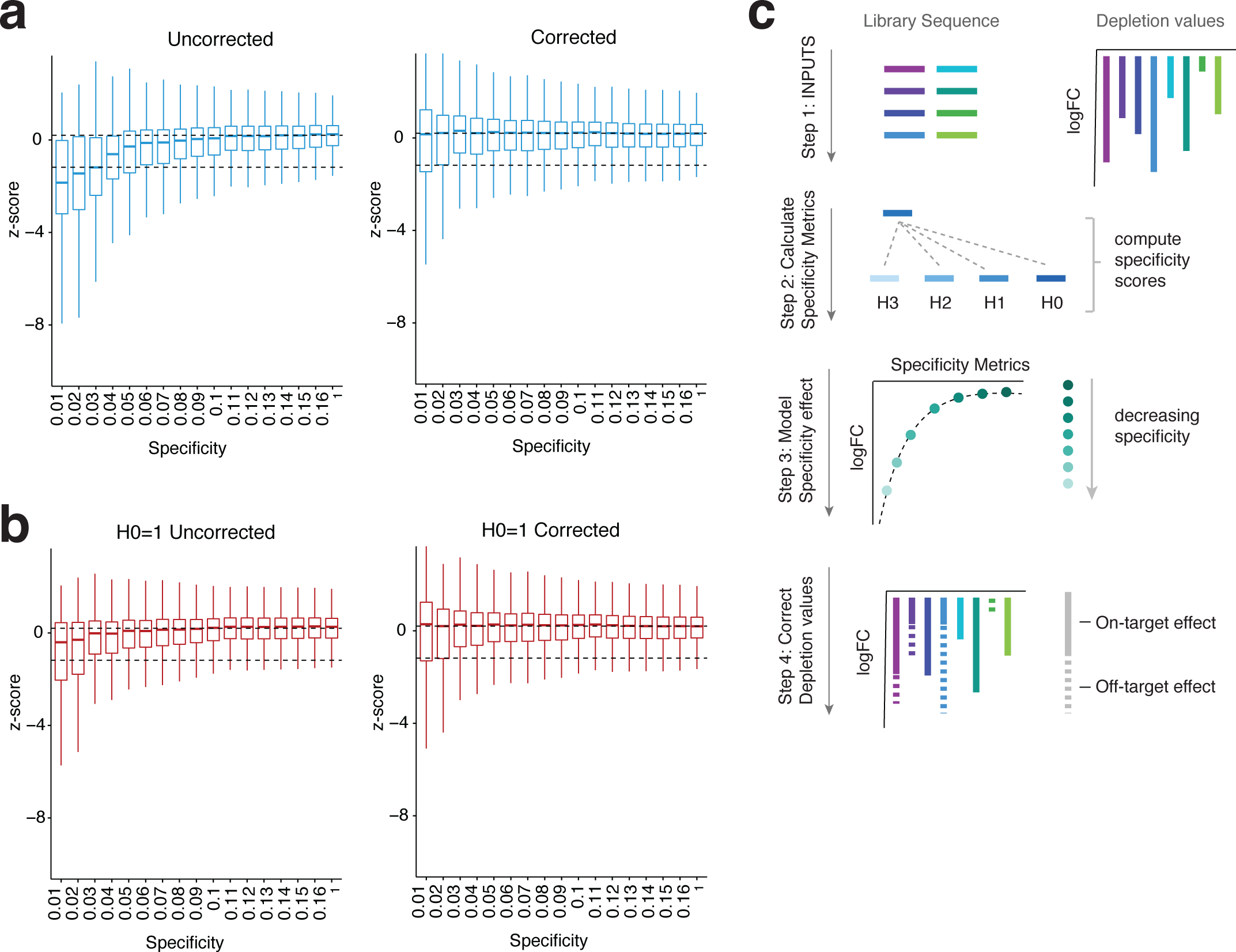
Computational correction of off-target mediated gRNA depletion from CRISPR-Cas9 essentiality screens. **(a)** Box plots of z-scores for gRNA log2-fold changes for all DepMap 19Q4 screens are shown across multiple Specificity bins before (left) and after (right) correction. Specificity values correspond to highest value of each bin. Dashed lines indicate mean of known non-essential (top) or essential (bottom) genes. **(b)** As in (a) but plotting only gRNAs that have a single perfect target site in the genome (H0=1; H0, hamming distance of 0). **(c)** Schematic representation of CSC. Briefly, the correction takes as input the gRNA sequences and corresponding depletion values. For each of the gRNAs, CSC enumerates all off-targets up to Hamming distance of 3 (H3), and computes its specificity score. CSC then uses the off-target information and depletion values to model the effect of off-targeting on gRNA abundance. Finally, it outputs both the corrected depletion values and the specificity metrics for each of the gRNAs. logFC, log fold change. Boxplots show minimum, maximum, median, first and third quartiles.

To determine the extent to which off-target mediated gRNA depletion acted as a confounder in the Achilles dataset, we calculate the Bayes Factor (BF) for each gene in individual screens using Bagel^15^. In this context, BF is an assessment of gene essentiality, with positive values indicating a gene is essential and negative values indicating a gene is non-essential. Gene Set Enrichment analysis (GSEA) showed that genes targeted by promiscuous gRNAs were significantly enriched in high BF values, particularly as the number of promiscuous gRNAs per gene increased or as the specificity of the gRNAs decreased (see **Supplementary Figure 1d-1f** for an example cell line). This suggests that, in line previous reports^2, 3^, off-targeting may contribute to false-positive dependencies even when multiple independent gRNAs per gene are present in the library.

To address this issue, we developed CRISPR Specificity Correction (CSC), a computational approach that removes the confounding effect of off-targeting from gRNA depletion (**Figure 1c**). CSC takes as inputs the sequence and depletion values of the gRNAs in a screen, and in a first step computes for each guide the number of target sites it has with zero (H0), one (H1), two (H2), or three (H3) mismatches. It also calculates GuideScan’s specificity score for each gRNA. CSC then uses these confounder metrics as covariates to model the contribution of off-targeting to gRNA depletion via a multivariate adaptive regression spline (see methods). Specificity-corrected depletion values for each guide are outputted with gRNA off-target information.

We applied CSC to all screens from the DepMap 19Q4 Achilles dataset. As predicted, this removed the correlation between guide specificity and its depletion (**Figure 1a, 1b**). More importantly, when inferring gene essentiality in each cell line using Bagel, correction of off-targeting by CSC significantly improved the recall of constitutive essential genes^16^, at a 5% false discovery rate (FDR) of constitutive non-essentials^16^ (**Figure 2**). This was true when looking at aggregate data for the entire data set (**Figure 2a**), as well as at the level of individual lineages or cell lines (**Figure 2b, 2c, Supplementary Figure 2**). Of note, CSC markedly outperformed the filtering of promiscuous gRNAs implemented by Project Achilles^6^ (**Figure 2**).

**Figure 2.**
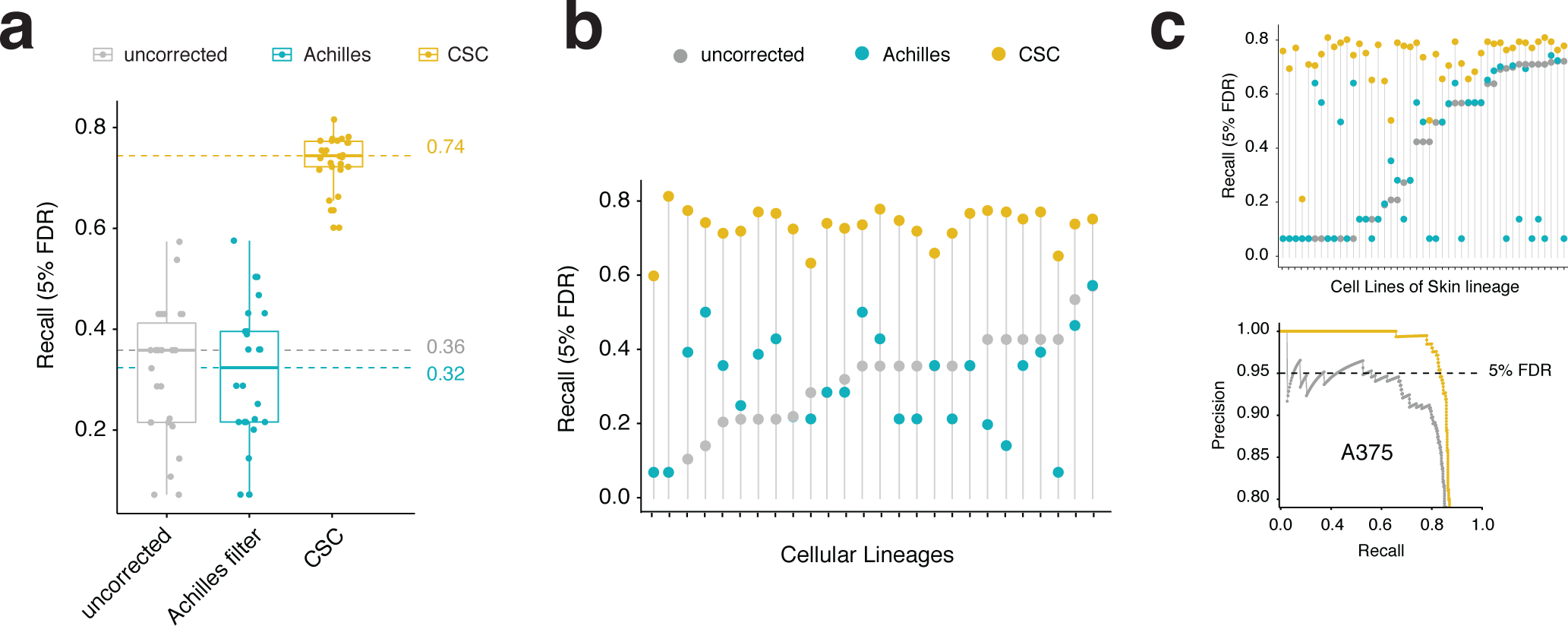
CSC improves the performance of CRISPR-Cas9 essentiality screens. **(a)** Boxplot showing recall values at 5% FDR for the 19Q4 DepMap dataset before correction (grey), with the Achilles filter (blue), or with the CSC correction (yellow). Each dot represents the median recall value of a lineage. Minimum, maximum, median, first and third quartiles are shown. **(b)** Recall values at 5% FDR for each lineage. **(c)** Top, example plot for a single lineage (skin). Bottom, example precision-recall plot for a single cell line (A375 melanoma cells). Lollipop graphs are plotted by increasing values of the uncorrected pipeline. The maximum recall value at 5% FDR was used for each comparison.

With the increased recall at 5% FDR, we observed a concomitant increase in the number of genes identified as essential (**Figure 3a**). In total, 12,444 genes scored as a dependency in at least one screen when CSC was implemented, compared with 5,831 and 6,018 genes for the uncorrected data or when off-targets were removed with the Achilles filter respectively. To attempt to discriminate which of these constituted true gene essentialities, we first examined the expression levels of genes inferred as essential by each pipeline. We found that those identified after off-target correction by CSC tended to be well expressed in the cell line in which they scored as essential (**Figure 3b**). In contrast, for each screen, genes scoring before, but not after, the correction tended to have significantly lower expression levels in that cell line. In fact, a substantial subset of these genes was not expressed at all, suggesting that they play no functional role in that context and thus represent false-positive hits. The same was true for genes identified after the implementation of the Achilles filter but not after CSC correction (**Figure 3b, Supplementary Figure 3a**).

**Figure 3.**
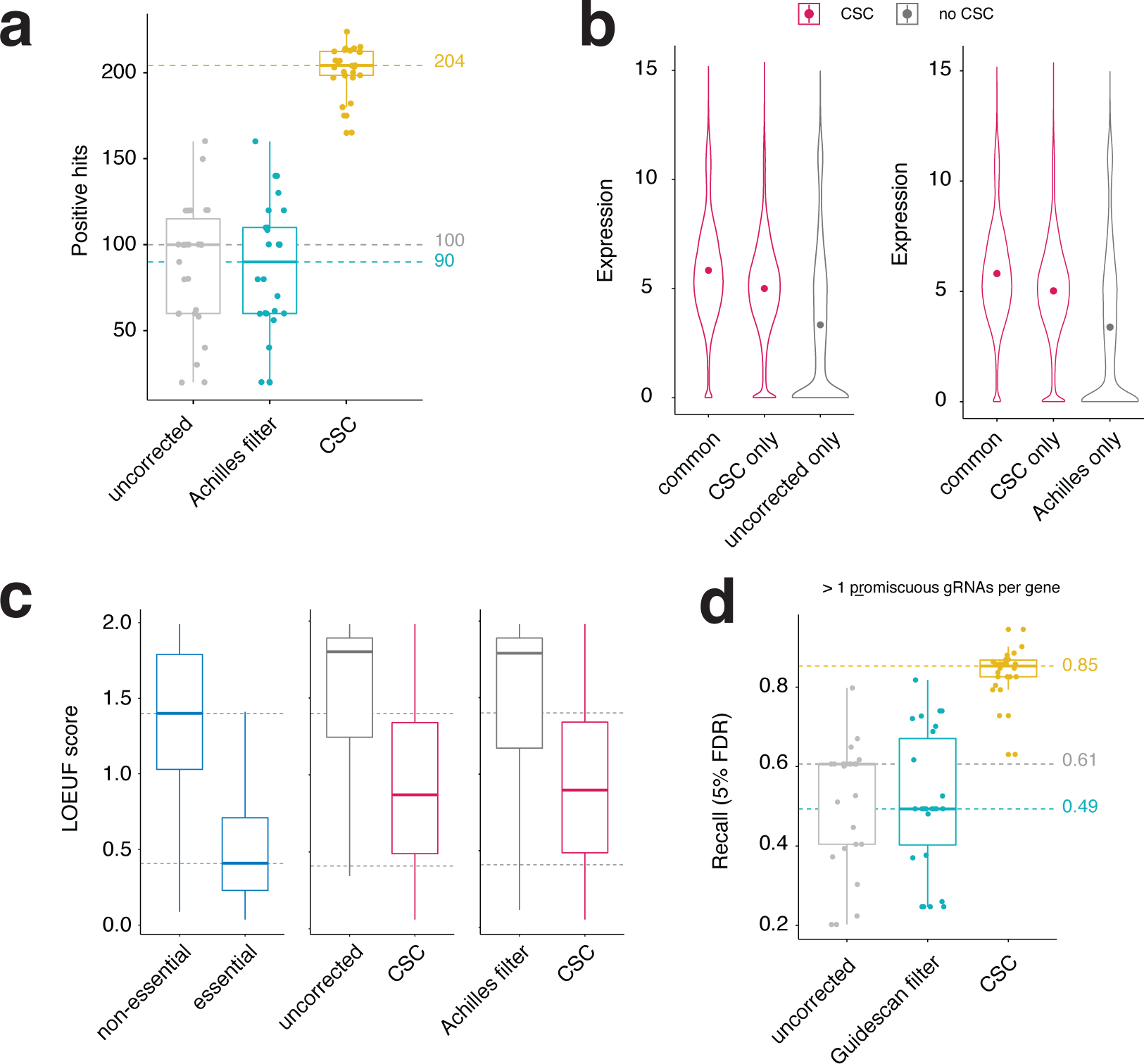
CSC improves the identification of true gene dependencies. **(a)** Boxplot showing number of positive hits at 5% FDR for the 19Q4 DepMap dataset before correction (grey), with the Achilles filter (blue), or with the CSC correction (yellow). Each dot represents the median number of dependencies in a lineage. **(b)** Violin plots showing the expression levels (log2(TPM+1)) of genes in the cell lines in which they were identified as dependencies. Left graph shows genes identified both before and after correction (common), only after correction (CSC only) or only before correction (uncorrected only). Right graph shows genes identified as dependencies both in pipelines that implement CSC and Achilles (common), or only in one of the pipelines (CSC only, Achilles only). Dot represents the mean value **(c)** Boxplots showing the LOEUF scores for genes identified as dependencies by only one of the pipelines. Exclusive dependencies are defined as genes that score in more than 15 cell lines in one pipeline and none in the other. LOEUF scores for constitutive essential and non-essential genes are shown on the left as a reference, and their median values are highlighted across all plots with a dashed line. **(d)** Recall values as in Figure 2a, but calculated using known constitutive essential and non-essential genes targeted by at least one promiscuous gRNA (H0 > 1). Boxplots show minimum, maximum, median, first and third quartiles.

Next, we looked for evidence of functional essentiality for genes identified as putative dependencies. We took advantage of the Genome Aggregation Database (gnomAD), which catalogs high-confidence predicted loss-of-function variants and uses these to classify human genes according to the mutational constrain they are under^17^. Specifically, the LOEUF score places genes along a spectrum of tolerance to inactivating mutations, where genes that play essential cellular roles, and therefore are under high mutational constraint, receive low LOEUF scores, while genes whose disruption has no impact on cell viability or organismal health and are therefore under low mutational constraint in the human population receive high scores (**Figure 3c**, left). We determined exclusive dependencies by selecting genes that scored in more than 15 screens in one pipeline but in no screens in the other. Genes within the CSC sets tended to have low LOEUF values, on average well below those attributed to constitutive non-essential genes (**Figure 3c**), suggesting that they may be under mutational constraint and therefore involved in essential functions. In contrast, genes that scored in more than 15 distinct screens only before the correction or only after removing promiscuous gRNAs through the Achilles filter but not in the CSC pipeline tended to have high LOEUF values, often above those of constitutive non-essentials suggesting that their inactivation is well tolerated in humans.

Taken together, the data presented above is consistent with the notion that genes identified after computational correction by CSC reflect true essentialities, and that CSC implementation minimizes the occurrence of false-positive hits. It also suggests that CSC outperforms the current filtering approach implemented by Project Achilles to deal with unspecific gRNAs (**Figure 3a-3c, Supplementary Figure 3a**). Because our initial analysis of the Achilles filter showed that it underestimated the promiscuity of the Avana library (**Supplementary Table 1**), we next set out to benchmark CSC against a filtering approach that accurately removes highly unspecific gRNAs. We reanalyzed all screens from the DepMap 19Q4 dataset after removing guide RNAs that had more than one perfect target site in the human genome (H0 > 1; GuideScan filter), and inferred gene essentiality as before using Bagel. This filtering significantly improved both the recall of constitutive essentials (**Supplementary Figure 3b**) and the number of positive hits (**Supplementary Figure 3c**) when compared to uncorrected or Achilles-filtered data. This suggests that the simple removal of highly promiscuous gRNAs can increase the sensitivity of Cas9 essentiality screens. Nevertheless, this approach still underperformed when compared to CSC (**Supplementary Figure 3b, 3c**) particularly as the number of promiscuous gRNAs per gene increased (**Figure 3d, Supplementary Figure 3d**). Indeed, when calculating precision and recall values for genes targeted by at least one highly promiscuous gRNA (H0 > 1) CSC still substantially increased recall (5% FDR) across all lineages of the dataset (**Figure 3d**) suggesting our computational approach is able to retrieve known gene dependencies even from highly promiscuous gRNA pools. In contrast, the simple removal of promiscuous gRNAs through the GuideScan filter led to decreased recall over this gene set when compared to the uncorrected data (**Figure 3d**), indicating that it is not a feasible approach for highly unspecific libraries.

In summary, we show that CSC is a significant improvement over existing approaches to deal with the confounding effect of unspecific gRNAs in CRISPR-Cas9 essentiality screens. We believe CSC will be a powerful aid to ongoing efforts to catalog genomic loci required for cellular fitness, particularly in the context of screens targeting highly repetitive genomic regions— such as non-coding regulatory elements—where gRNA filtering is often not feasible^2^. To facilitate its incorporation into existing CRISPR analysis pipelines, we make the software freely available as a python package at https://bitbucket.org/arp2012/csc_public/src/master/

## METHODS

### Screening data

Raw read counts of CRISPR viability screens (Broad DepMap project 19Q4) performed with the Avana library were downloaded from the DepMap project data repository (https://figshare.com/articles/DepMap_19Q4_Public/11384241/2).

### Guide RNA preprocessing

Guide RNAs with less than 30 reads in the initial plasmid counts were removed. Counts for all screen replicates and corresponding plasmid library were adjusted by median-ratio normalization to adjust for the effect of library sizes and read count distributions. Finally, for each screen, log_2_-fold changes for individual gRNAs were calculated between the initial plasmid library counts and the post-screen counts for each replicate experiment. The mean log_2_-fold changes between replicates was used as the final log_2_-fold changes value for each gRNA.

### Off-target data

We downloaded the sequence and annotation data for the hg38 assembly of the human genome from the UCSC database^18^ and used it to construct a retrieval tree (trie) consisting of all possible 20mer Cas9 gRNA target sites in the human genome with the GuideScan software^13^. In contrast to the original trie^13^, this retrieval tree was constructed without inclusion of alternative chromosome data, so that they did not artificially inflate the enumeration of off-targets. To determine the mismatch neighborhood for each gRNA in the Avana library, we traversed their sequences through the trie to exhaustively determine all neighbors up to and including Hamming and Levenshtein distances of 3. Specificity scores for each gRNA were computed using Hamming distance neighbors using our previously described strategy^13^, which incorporates Guidescan’s ability to faithfully enumerate all potential target sites up to a specified number of mismatches, with CFD score’s prediction of how likely each of those sites is to be cut^14^. All these metrics are used as covariates (*x*) in the CSC model (see below).

### Multivariate Adaptive Regression Splines

We developed a model that assumes that the measured depletion value (*D*) of a gRNA (*i*) in any individual screen is the sum of gene-knockout effects (*G*_*i*_) and off-target effects (*O*_*i*_)

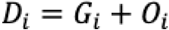

To estimate *O*_*i*_, we use Multivariate Adaptive Regression Splines^19^ (EARTH) which can model non-linearities in the data as well as interactions between variables. The model takes the form of the following equation

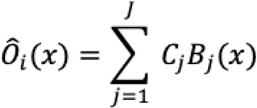

where the estimated contribution of off-targeting to gRNA depletion (*Ô*_*i*_) can be approximated by the weighted sum of *J* basis functions *B*_*j*_ derived from the model predictor variables (*x*). *C*_*j*_ are coefficients of expansion whose values are jointly adjusted to give the best fit to the data. The basis functions *B*_*j*_ can take the form of i) a constant 1, which represents the intercept of the model; ii) a hinge function derived from a predictor variable, or iii) a product of two or more hinge functions each derived from different predictors to capture their interaction.

#### Model Training and Pruning

The model starts with the intercept term (, with intercept at). It then iteratively adds new basis functions in the form of hinge functions or products of hinge functions. At each step, the new terms are selected and added into the model as to minimize the sum of squared error using ordinary least squares method. This forward pass proceeds until the residual error consistently falls below the stopping threshold (minimal change in mean squared error (MSE) with additional terms). To prevent over-fitting and improve generalization, the forward pass model undergoes a backwards pass, where model terms are removed in a stepwise manner with subsequent reassessment for increases in the sum of squared error obtained in this sub-model. Selection for the optimal sub-model is done using generalized cross-validation (GCV) which optimizes tradeoff between bias and variance. The model with the lowest GCV is selected as the optimal model. An example file detailing model metrics for a DepMap screen is provided as **Supplementary File 1**.

In the development of this model training data consists of 90% of all input data; test data consists of the remaining 10%. Test error was assessed as root mean squared error between the predicted and actual values of test data.

### CSC Software and implementation

The CSC was packaged in Python (version 2.7.15) with Avana, Brunello, and GeckoV1 and GeckoV2 libraries as package data. Pickle files for hg38 and mm10 genomes are also provided in a repository, to allow CSC to be implemented for any custom human or mouse libraries. We also provide a Docker image. All these files are freely available to download from our bitbucket repository (see Code Availability).

### Alternative approaches for gRNA off-targeting correction

To benchmark CSC, we compared its performance against the current approach of filtering out gRNAs suspected of off-target activity. We tested two distinct filtering strategies. The first, (which we refer to as Achilles filter) is based on the strategy adopted by Project Achilles to remove the confounding effect of off-targeting from the CRISPR essentiality screens performed with the Avana library. Information about this filter (which can be downloaded from the DepMap data repository as “Achilles_dropped_guides.csv”) is provided here as part of Supplementary Table 1 (columns 8 and 9). The Achilles filter list was generated from runs of CERES, and includes guides flagged for potential off-target activity by CERES based on being the sole efficacious gRNA for a gene receiving a label of “guide_dropped_by_ceres”. In addition, this file enumerates the estimated number of perfect matches for each guide in the column “Achilles n_alignments”. These alignments are performed with Bowtie against the 20-nucleotide long sequence of each gRNA and subsequently filtered for the presence of PAM sequence motif in the form of NGG^6^. gRNAs that have no perfect alignment to hg38 or that are found to have more than one perfect target site through this method are dropped from the analysis and flagged as “not_aligned” or “in_ dropped_guides” respectively.

As discussed in Supplementary Note 1, and shown Supplementary Table 1, the filtering list that is generated through the method described above significantly underestimates the number of promiscuous gRNAs in the Avana library. In fact, estimation of perfect target sites by this approach only surpasses that of the GuideScan retrieval trie algorithm in 8 cases:

| gRNA ID | H0 (GuideScan) | Achilles n_alignments |
| --- | --- | --- |
| 3560 | 2 | 3 |
| 23187 | 9 | 12 |
| 23692 | 4 | 5 |
| 55070 | 1 | 4 |
| 67756 | 4 | 6 |
| 69883 | 1 | 2 |
| 93689 | 1 | 2 |
| 93814 | 9 | 11 |

Manual curation of each of these shows that the additional sites identified by Bowtie but not GuideScan were not adjacent to PAM motifs, and therefore do not represent potential target sites for the guide RNA in question.

Because the Achilles filter underestimated the number of promiscuous gRNAs in the library, we also included an additional control pipeline in which only gRNAs with a single perfect target site, as identified by GuideScan, were kept. The identity of these guides is described in Supplementary Table 1.

Each of these three approaches (CSC, Achilles filter, GuideScan filter) was integrated within otherwise identical analysis pipelines and compared against a pipeline in which no off-target management was performed.

### Precision-recall analysis

To compare the performance of the methods described above, we generated precision-recall curves for all screens after being processed through each of the four analysis pipelines, using the set of constitutive essential and non-essential genes defined in *Hart et al*.^16^, as references. Curves were generated based on ordered BF values (see below). The BF value corresponding to 95% of precision (meaning the value for which at least 95% of genes are known essentials) was taken as the 5% False Discovery Rate threshold (FDR = 1 - precision). The percentage of reference essential genes identified as essential at that BF threshold was taken as the recall value at 5% FDR. In cases where multiple threshold values had a precision of 95%, that corresponding to the highest recall value was used.

### Inference of gene essentiality

The Bagel software (version 0.91)^15^ was used to infer gene essentiality based on log_2_-fold changes of gRNAs for each gene. This software uses a supervised learning method which implements Bayesian statistics and outputs for each gene a Bayes Factor (BF) value based on the likelihood that the observed fold-changes of the gRNAs that target it were drawn from reference essential or non-essential distributions^15^. For each screen, essential genes at 5% FDR we identified by selecting those with BF values above the threshold identified in the precision-recall analysis.

### Gene expression data

RNA-seq TPM gene expression data (log_2_-transformed using a pseudo-count of 1) for protein coding genes was downloaded from the DepMap project data repository (https://figshare.com/articles/DepMap_19Q4_Public/11384241/2).

### Code availability

Scripts for off-target enumeration and CSC implementation are freely available at our Bitbucket repository (https://bitbucket.org/arp2012/csc_public/src/master/).

## Supporting information

Comparison of the off-target description for the Avana library by Achilles filter or GuideScan

Example metrics file generated by the CSC software

## ACKNOWLEDGMENTS

We would like to thank all members of the Vidigal lab for critical comments on the manuscript. We thank Ralph Garippa and Michael Lichten for suggesting the development of CSC. Eros Denchi, Luke E Dow, Lee Zamparo, and Ram Kannan provided critical feedback on the manuscript. ARP thanks Olaf Andersen for mentorship and support throughout his professional training. This work utilized the computational resources of the NIH HPC Biowulf cluster (http://hpc.nih.gov) and the UCSF Wynton cluster (https://wynton.ucsf.edu), and was supported by the Intramural Research Program of the National Institutes of Health (NIH) and a FLEX grant from the Center for Cancer Research (JAV).

## AUTHOR CONTRIBUTIONS

A.R.P. and J.A.V. conceived and designed the study. A.R.P. wrote and implemented all software. A.R.P., L.S., and J.A.V. processed, managed, and analyzed the data. R.K.P. assisted with computational analysis. J.A.V, A.R.P., and L.S. wrote and/or revised the manuscript with assistance from R.K.P.. J.A.V. supervised the study.

## Supplementary Note 1

The confounding effect of off-targeting has been well documented in CRISPR assays and, over the past several years, numerous research efforts have aimed at defining the tolerance of Cas9 to mismatches^14, 20, 21^. As a result, we now have a fairly comprehensive understanding of how the number, position, and type of mismatches interfere with both Cas9 binding^22^ and endonucleolytic cleavage^14, 20-22^, leading to the development of scores such as CFD^14^ and Elevation^21^ which describe how likely a potential off-target sequence is to be cleaved based on its similarity to the gRNA.

Despite this wealth of knowledge, the strategies employed to identify the genomic sequences that constitute potential off-target loci—on which CFD and Elevation scores are calculated—can be inaccurate^13^. Specifically, identification of potential sites of off-target cleavage is often achieved through the use of short-read aligners. However, aligners such as Bowtie^23^, Bowtie2^24^, STAR^25^, and even BLAT^26^ have a trade-off between speed and exhaustive read-matching, leading to the truncation of a search if an effort limit is exceeded^23-26^. This feature allows tools to process queries quickly, an essential criterion when working with large datasets. Yet, in the context of CRISPR off-target search it means that exhaustive identification of off-target sites is not guaranteed particularly for highly promiscuous gRNAs. As a result, many gRNA design algorithms that incorporate aligners in their off-target search method under-enumerate potential off-target loci for individual gRNAs^13^. We find that, contrary to what has been previously reported^14^, this holds true for sites that have perfect matches to the gRNA^13^— as exemplified below for the most promiscuous gRNA in the Avana library—and for tools commonly thought to perform comprehensive off-target searches^27^:

**Supplementary Table 2:**
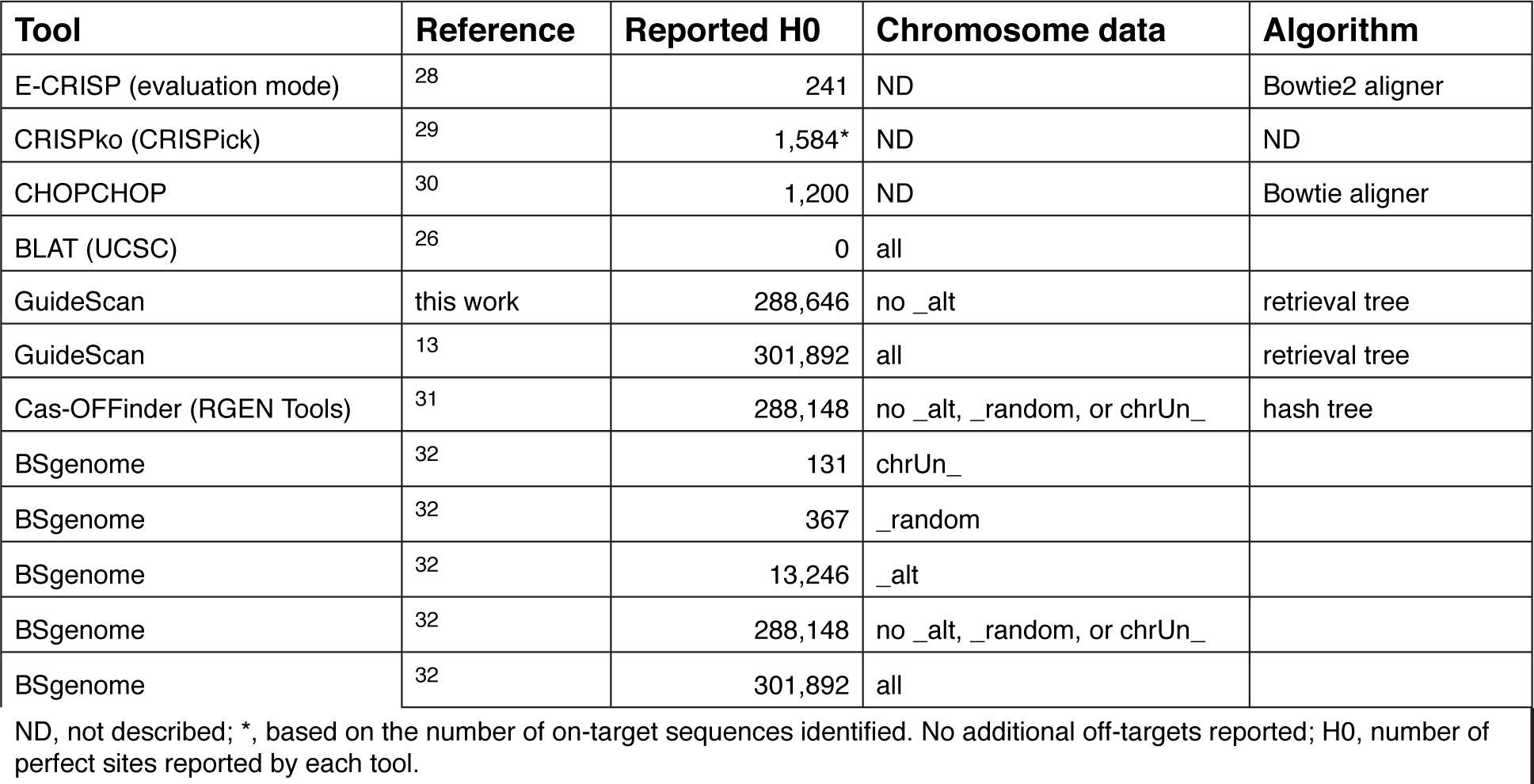
Comparison of off-target enumerations by commonly used Tools for the most promiscuous gRNA in the Avana library.

In contrast, both GuideScan^13^ and Cas-OFFinder^31^—which do not utilize aligners in their search algorithm— accurately enumerate potential gRNA off-target loci, yielding identical counts to those of an exhaustive search performed with the BSgenome R package^32^.

Project Achilles performs alignments using Bowtie as part of their off-target search method^6^, and that likely accounts for the underestimation of gRNA promiscuity in the Avana library in the Achilles off-target filter (see also **Supplementary Table 1**).

**Supplementary Figure 1.**
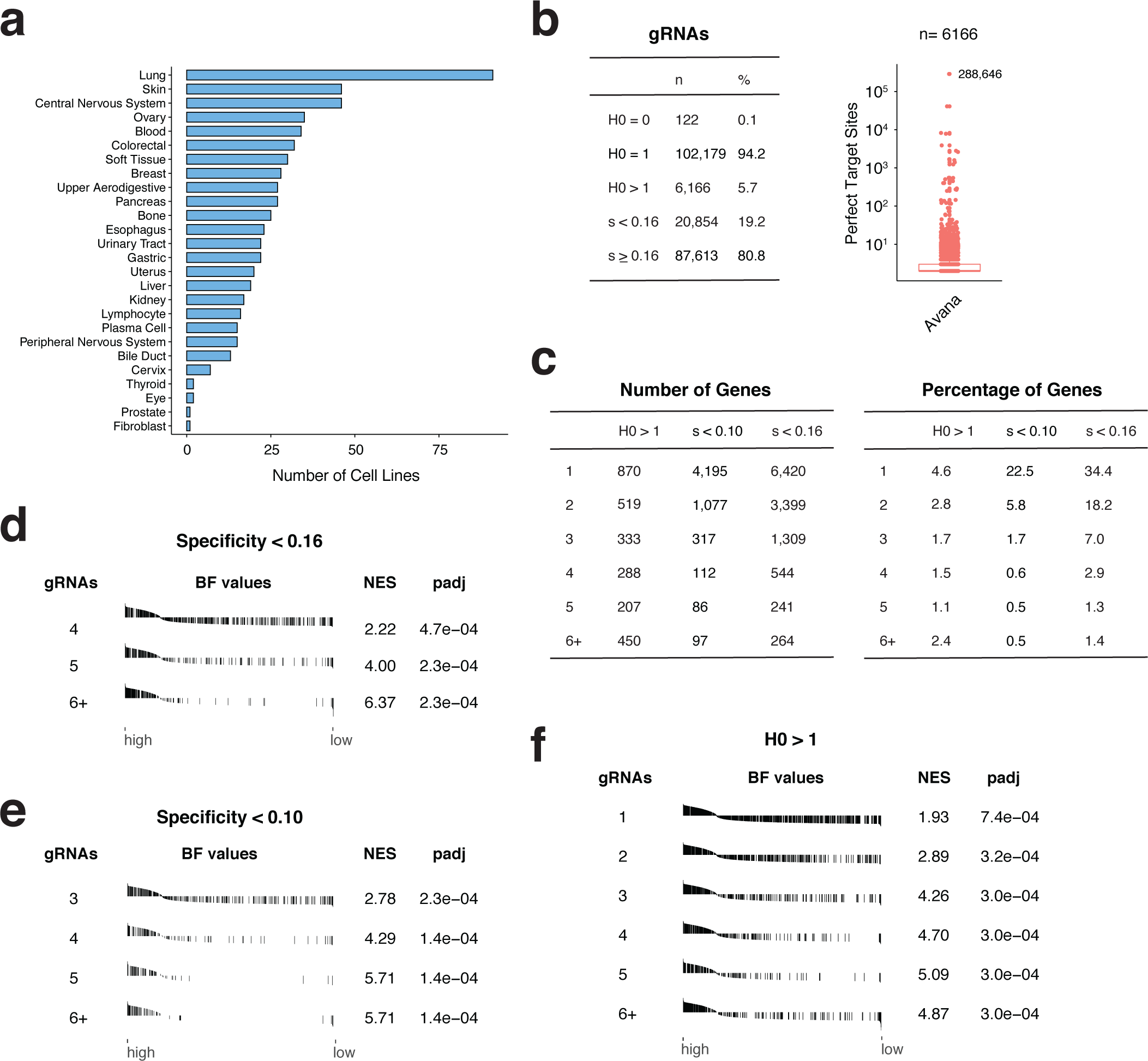
Analysis of the 19Q4 Avana dataset from the DepMap initiative. **(a)** Number of screened cell lines in each of the 26 lineages represented in the dataset. **(b)** Left, number and percentage of gRNAs in the Avana library that have 0, 1, or more than 1 perfect targets (H0) in the human genome (hg38 assembly) or that have specificities (s) lower or equal/higer than 0.16. Right, number of perfect target sites for gRNAs with H0 > 1. The gRNA in the Avana library with the highest number of perfect targets sites is highlighted. Boxplot shows minimum, maximum, median, first and third quartiles. **(c)** Summary of the number (left) and percentage (right) of genes targeted by increasing number of promiscuous gRNAs (1-6+). Promiscuity is defined at three distinct thresholds (H0 > 1, s < 0.10, s < 0.16). **(d-f)** Gene set enrichment analysis for the A375 melanoma cell line screen showing that genes targeted by promiscuous gRNAs (defined as specificity < 0.16 (d), specificity < 0.10 (e), or presence of multiple perfect targets (H0 > 1, f)) are significantly enriched in high BF values. Each line of the plot represents a gene in the set, ordered by decreasing BF values. Gene sets were defined by number of promiscuous gRNAs targeting a gene. Only significant gene sets are shown. NES, normalized enrichment score. padj, p value adjusted for multiple testing.

**Supplementary Figure 2.**
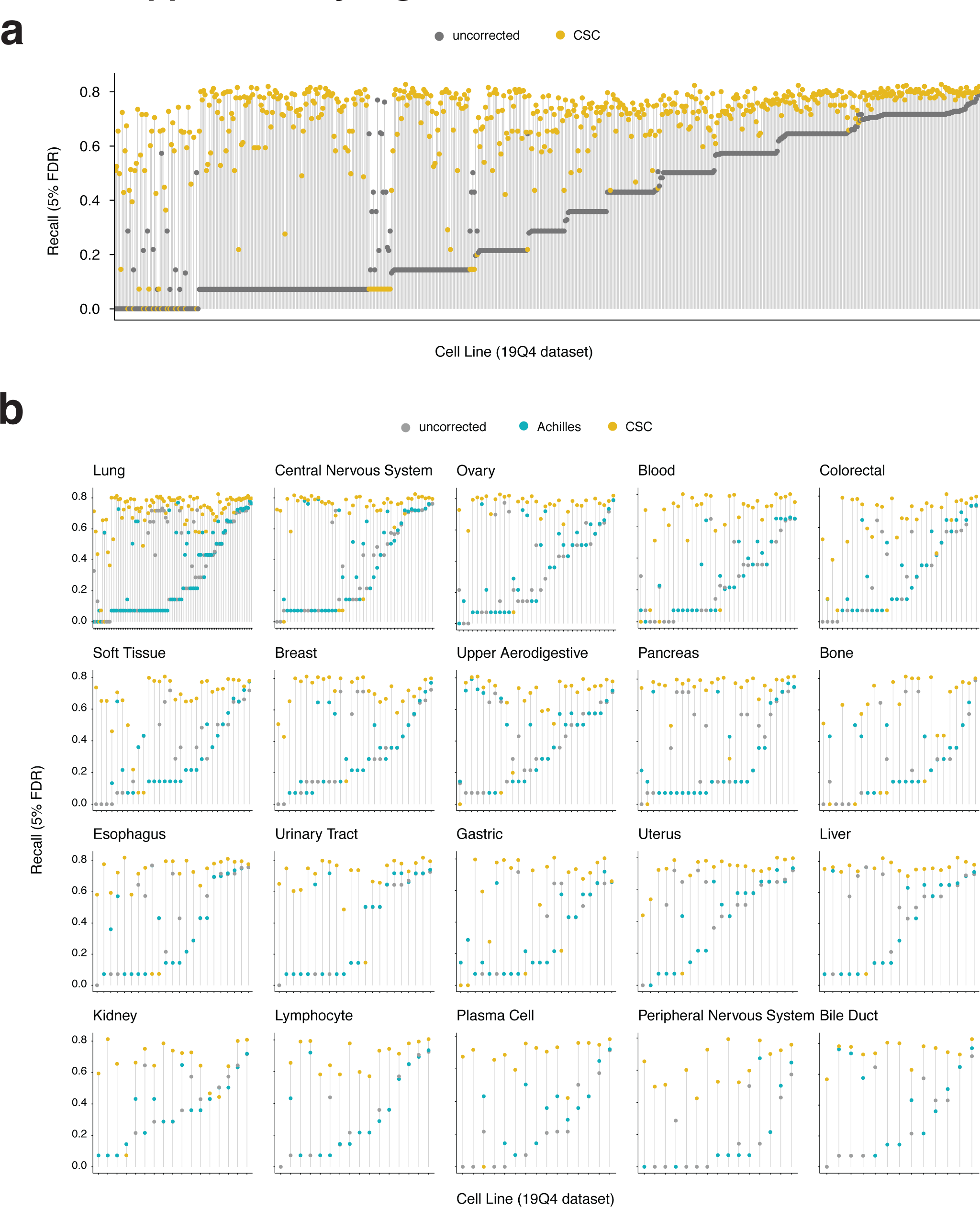
CSC improves the performance of CRISPR-Cas9 essentiality screens. **(a)** Recall values at 5% FDR for each cell line in the 19Q4 dataset before (grey) or after correction (yellow). **(b)** Recall values before and after correction, plotted by lineage. Only lineages with more than 10 screens are shown.

**Supplementary Figure 3.**
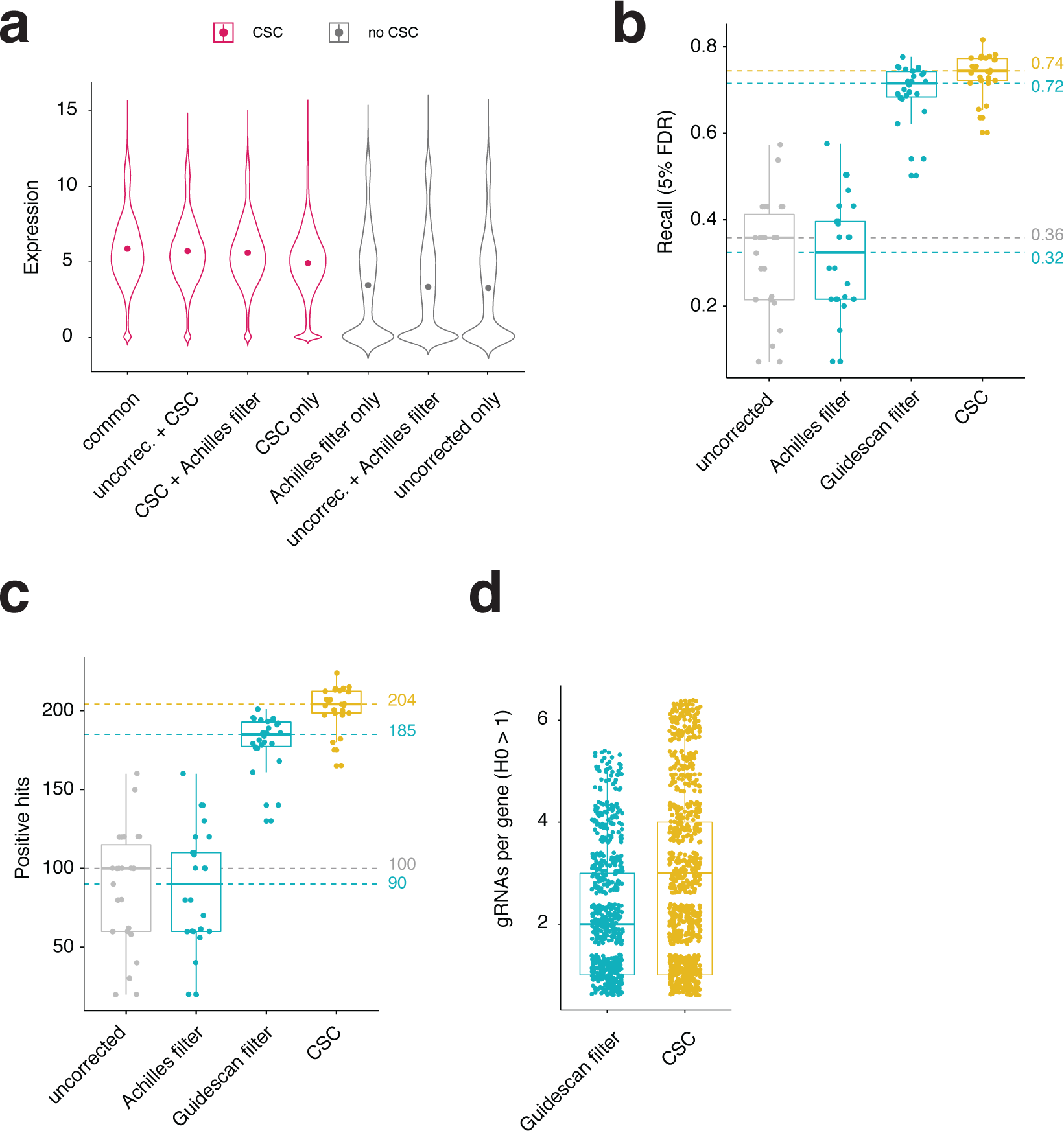
Performance of CSC against promiscuous gRNA filtering. **(a)** Violin plots showing the expression levels (log2(TPM+1)) of genes in the cell lines in which they were identified as dependencies. The common dataset shows genes identified by all three pipelines in a subset of cell lines. Dot represents the mean value. **(b)** Recall values at 5% FDR for the 19Q4 DepMap dataset before correction (grey), with gRNA filtering (blue), or with the CSC correction (yellow). Each dot represents the median recall value of a lineage. **(c)** as in (b) but showing the number of genes identified as essential by each pipeline. **(d)** Boxplot showing the number of promiscuous gRNAs (H0 > 1) targeting genes identified as essential by each pipeline. Each dot represents a gene. Only genes targeted by at least one promiscuous gRNA included. Boxplots show minimum, maximum, median, first and third quartiles.

